# *Thy1* (CD90) expression is regulated by DNA methylation during adipogenesis

**DOI:** 10.1101/365809

**Authors:** E’Lissa M. Flores, Collynn F. Woeller, Megan L. Falsetta, Martha Susiarjo, Richard P. Phipps

**Author notes:** Corresponding author, Corresponding author (to whom all correspondence should be addressed): Collynn Woeller, Ph.D. Research Associate Professor University of Rochester Department of Environmental Medicine 601 Elmwood Ave, Box 850, Room 3-11001 Rochester, NY 14642. Work funded by F31ES027767, TL1-TR000096, ES001247, and ES0023032.

## Abstract

The obesity epidemic is developing into the most costly health problem facing the world. Obesity, characterized by excessive adipogenesis and enlarged adipocytes, promotes morbidities such as diabetes, cardiovascular disease and cancer. Regulation of adipogenesis is critical to our understanding of how fat cell formation causes obesity and associated health problems. Thy1 (also called CD90), a widely used stem cell marker, blocks adipogenesis and reduces lipid accumulation. Thy1 knockout-mice are prone to diet-induced obesity. While the importance of Thy1 in adipogenesis and obesity is now evident, how its expression is regulated is not. We hypothesized that DNA methylation plays a role in promoting adipogenesis and affects *Thy1* expression. Using the methylation inhibitor 5-aza-2’-deoxycytidine (5-aza-dC), we investigated whether DNA methylation alters *Thy1* expression during adipogenesis in both mouse 3T3-L1 pre-adipocytes and mouse mesenchymal stem cells. Thy1 protein and mRNA levels were decreased dramatically during adipogenesis. However, 5-aza-dC treatment prevented this phenomenon. Pyrosequencing analysis shows that the CpG sites at the *Thy1* locus are methylated during adipogenesis. These new findings highlight the potential role of *Thy1* and DNA methylation in adipogenesis and obesity.

## Introduction

Obesity rates have risen markedly in the last 30 years. More than 700 million people worldwide are clinically obese^1,2^. Obesity promotes type 2 diabetes, fatty liver disease, and cardiovascular disease, and is linked with certain cancers^3-5^. Health care costs associated with obesity and its comorbidities are enormous and will continue to rise as currents trends continue and the population ages^6^. Thus, a further understanding of obesity and its underlying mechanisms and causes are urgently needed.

Obesity results from a positive energy balance when more calories are consumed than are used. The surplus energy is packaged into lipid-based storage molecules and sent to fat storage cells called adipocytes. In obesity, there is both an increase in adipocyte size and an increase in adipocyte number to accommodate the lipid^7^. Adipocytes are formed during the process of adipogenesis and arise from stem cells, fibroblasts, or other progenitor cells when appropriately programmed^8^. Adipogenesis is a highly regulated process that requires the activation of several key signaling pathways, including STAT5 and Fyn and activation of the transcription factors PPARγ and C/EBPα. Numerous genes involved in fatty acid transport and storage, such as fatty acid binding protein 4 (Fabp4) are induced during adipogenesis to promote lipid accumulation in adipocytes.

Several proteins including Pref-1, Wnt, TGFβ, and Thy1 (formally called CD90) have been shown to inhibit adipogenesis by blocking pro-adipogenic signaling^9-13^. Our recent study showed that Thy1 blocked the activity of the Src family kinase, Fyn, in pre-adipocytes^14^. Thy1 mediated inhibition of Fyn activity prevented adipocyte formation. Interestingly, while pre-adipocytes expressed high levels of Thy1, its expression was lost during adipogenesis, and mature adipocytes expressed almost no Thy1. Thy1 is a member of the immunoglobulin supergene family and is a glycophosphatidyl inositol linked surface protein. While Thy1 is expressed on pre-adipocytes and subsets of fibroblasts, neurons, and stem cells, little is known about how its expression is controlled. We recently showed that Thy1 levels can be regulated by microRNAs. Specifically, the miR-103/107 family of miRNAs can target Thy1 mRNA and reduce its expression^15^. Furthermore, the *Thy1* gene contains several CpG rich elements termed CpG islands, which are hotspots for cytosine methylation and gene regulation^16,17^. To date, there have been no reports studying DNA methylation of *Thy1* during adipogenesis. However, *Thy1* methylation has been studied in context of fibrosis and in T cells, where they show an increase in *Thy1* methylation correlated with a decrease in *Thy1* expression^18-20^. Therefore, we investigated the same CG rich region within intron 1 (termed Thy1-CGI1), which is part of the promoter^17,20-23^.

While changes in DNA methylation patterns at the *Thy1* locus have not been characterized in the context of adipogenesis, recent reports have shown that DNA methylation changes are an integral part of adipocyte formation^24,25^. In the most widely used and well accepted model of adipogenesis, the murine 3T3-L1 pre-adipocyte line, global DNA methylation has been shown to increase during adipogenesis^26,27^. Interestingly, addition of the DNA methylation inhibitor 5-aza-2’-deoxycytidine (5-aza-dC) at the onset of adipogenic differentiation can completely block adipocyte formation, suggesting that DNA methylation changes are essential for adipogenesis^27,28^. Specific changes in regional DNA methylation are also critical for adipogenesis, and blocking these patterns can alter the differentiation pathway of precursor cells towards osteoblastogenesis rather than adipogenesis^24,29,30^. For example, when blocking DNA methylation using 5-aza-dC in 3T3-L1 cells, Wnt10a expression increases through the hypomethylation of several of its CpG sites. The increase in Wnt10a expression helps steer the differentiation program towards osteoblastogenesis and away from adipogenesis^30^.

Because Thy1 protein and mRNA levels are rapidly suppressed during adipogenic differentiation, we hypothesized that its reduced expression is linked to hypermethylation of the *Thy1* locus. Excitingly, we are the first to show methylation sensitive pyrosequencing data of the Thy1-CGI1 region and report herein that DNA methylation is one of the essential regulators of *Thy1* expression during adipogenesis.

## Results

### Reduced Thy1 expression during adipogenesis is partially attenuated by 5-aza-dC in bone marrow-derived mouse mesenchymal stem cells

Previously, we demonstrated that pre-adipocytes and mesenchymal stem cells must down-regulate *Thy1* expression to differentiate into adipocytes^14^. To determine if DNA methylation regulates *Thy1* expression during adipogenesis, we first pharmacologically inhibited DNA methyltransferase in bone-marrow derived mouse mesenchymal stem cells (mMSCs) before differentiation into adipocytes^28^. mMSCs were treated daily with either DMSO (vehicle) or the DNA methyltransferase inhibitor, 5-aza-dC for 12 days and were concurrently treated with either media alone or with the adipogenic cocktail medium (ACT), starting at day 3 (Fig. 1A). mMSCs typically differentiate into adipocytes in 2 weeks when exposed to ACT^31^. Therefore, we examined an intermediate time point to examine Thy1 levels. As expected, when given ACT, Thy1 levels decreased while fatty acid binding protein 4 (Fabp4) levels were increased compared to media alone (Fig. 1B-D); Fabp4 is highly expressed by adipocytes and serves as an adipogenic marker. However, inhibiting DNA methylation with 5-aza-dC increased Thy1 protein and mRNA levels versus media alone and attenuated the down-regulation of Thy1 in ACT samples (Fig. 1B-C). We also determined that protein and mRNA levels of Fabp4, a protein highly expressed by adipocytes that serves as an adipogenic marker, were decreased in mMSCs treated with 5-aza-dC (Fig. 1B,D). Immunofluorescence staining revealed that compared to cells not treated with ACT, 5-aza-dC-treated mMSCs had increased surface expression of Thy1 and decreased Fabp4 levels (Fig. 1E). These results indicate that inhibition of DNA methylation impairs adipogenesis.

**Figure 1:**
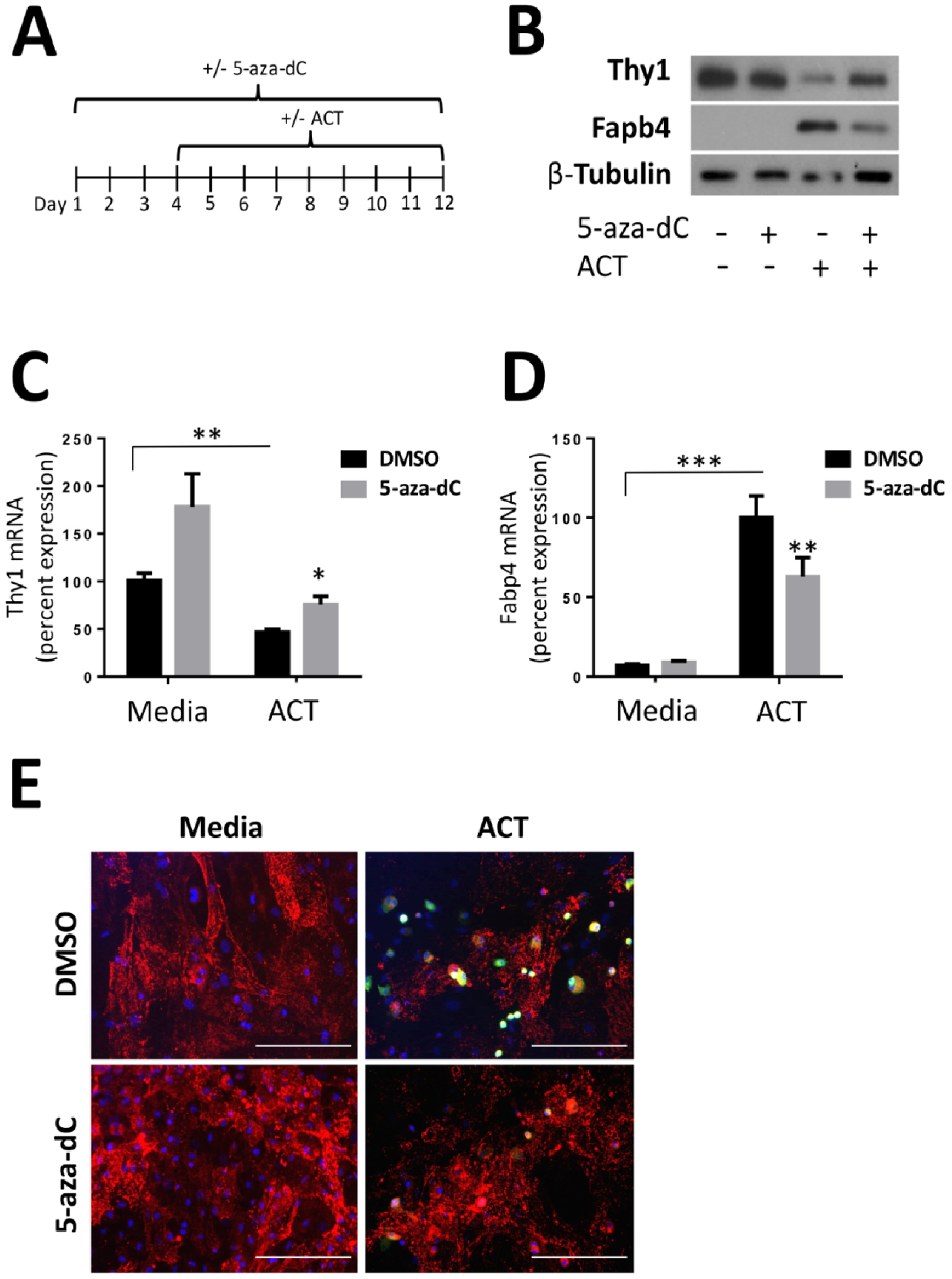
The reduction of *Thy1* expression during adipogenesis is attenuated by 5-aza-dC in mouse mesenchymal stem cells. mMSCs were treated daily with either DMSO or 5-aza-dC for 12 days continually with basal media alone or with the introduction of the adipogenic cocktail (ACT) starting at day 3 A) Timeline diagram of treatment for mMSCs. B) Western blot shows treatment with 5-aza-dC increases Thy1 protein levels and decreases protein levels of the adipogenic marker, Fabp4 versus cells only receiving the adipogenic cocktail (ACT). C-D) RT-qPCR results show that treatment with 5-aza-dC significantly increases Thy1 mRNA and decreases Fabp4 mRNA levels. Relative percentages were normalized to Media DMSO for Thy1 mRNA and ACT DMSO for Fabp4 mRNA levels. E) Immunofluorescent images depict reduced FAPB4 (Green) expression and increased Thy1 (Red) expression in cells receiving 5-aza-dC versus those receiving ACT alone. Cell nuclei are stained with DAPI and depicted in blue. Scale bars in white represent 200μm. *p<0.5, **p<.01, ***p<.001.

### Reduced Thy1 expression, which is necessary for adipogenesis, is prevented by inhibition of global DNA methylation in 3T3-L1 cells

Primary mesenchymal stem cells are heterogeneous with only some cells fully differentiating into adipocytes^32^. Thus, we next used the well-established pre-adipocyte murine cell line, 3T3-L1. Previously, we showed that fully differentiated adipocytes treated with ACT no longer expressed Thy1 protein and mRNA^33^. To observe changes in Thy1 expression, we examined an intermediate point in adipogenesis. At only four days of treatment with ACT the cells are only partially differentiated, and Fabp4 expression increases. Cells were treated daily for 7 days with either DMSO or 5-aza-dC and starting at day 3 were either given ACT or continued with media alone (Fig. 2A). As expected, Thy1 decreased at both the protein and mRNA level in cells given ACT compared to media alone (Fig 2B-C). However, we show that treatment with 5-aza-dC prevents the decrease in Thy1 mRNA levels in ACT-treated cells, while media alone with 5-aza-dC causes no significant changes (Fig 2C). 5-aza-dC also caused a decrease in Fabp4 levels in ACT-treated cells, indicative of decreased adipogenesis (Fig 2B,D). Inhibiting global methylation blunts adipogenesis, which is reflected by the decreased levels of Fabp4 expression and in part sustains Thy1 levels.

**Figure 2:**
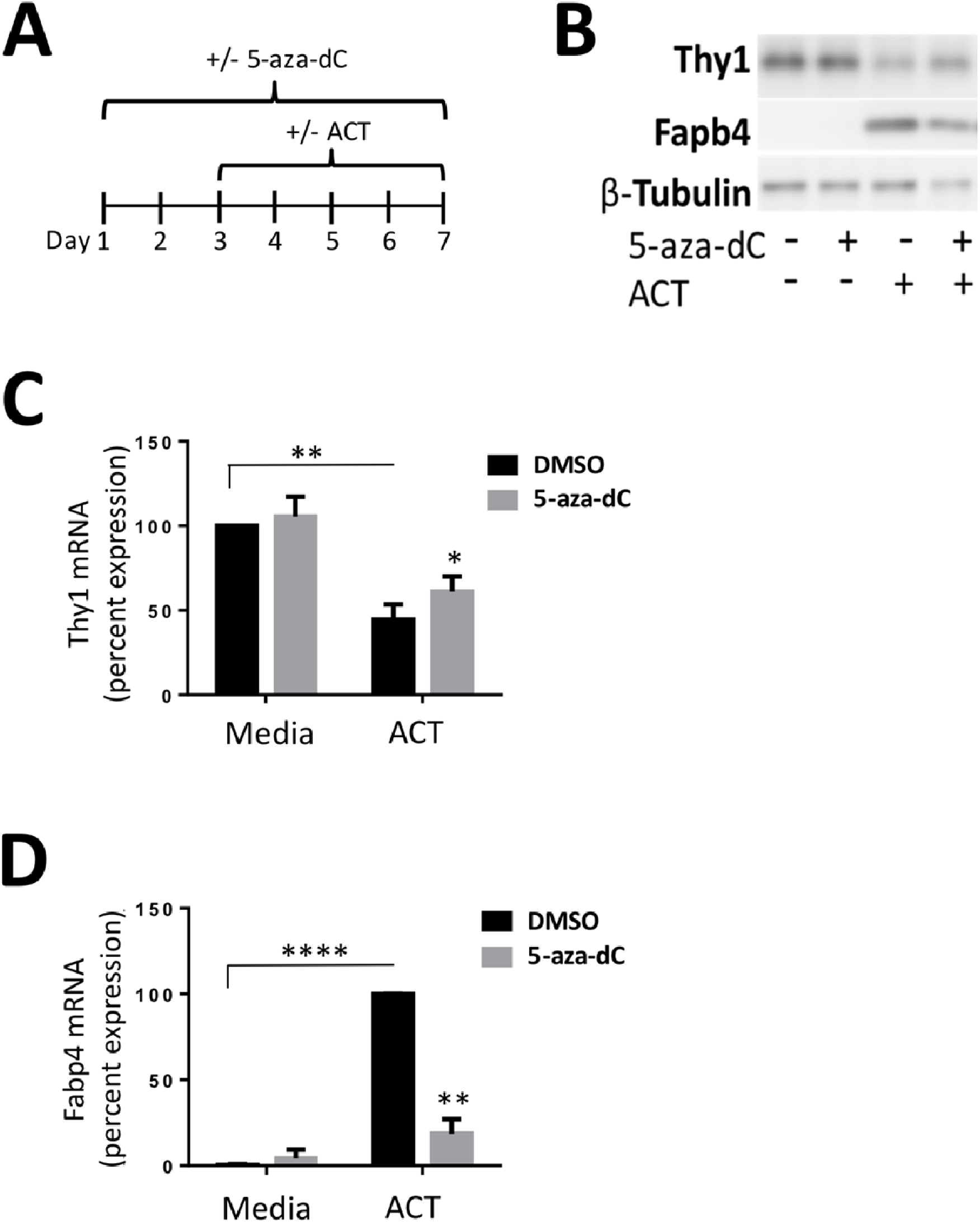
Inhibiting methylation increases *Thy1* expression in 3T3-L1 during adipogenesis. 3T3-L1 cells were treated daily with either DMSO or 5-aza-dC for 7 days continually with basal media alone or with the introduction of the adipogenic cocktail (ACT) starting at day 3. A) Timeline diagram of treatment for 3T3-L1s. B) Western blot showing treatment with 5-aza-dC resulted in an increase in Thy1 total protein levels in both media alone and adipogenic cocktail exposed samples. ACT samples also had a decrease in Fabp4 protein levels when treated with 5-aza-dC, shown by a representative western blot. C-D) RT-qPCR shows treatment with 5-aza-dC increases Thy1 mRNA and decreases Fabp4 mRNA levels in ACT samples. Relative percentages were normalized to 100% to media DMSO for Thy1 mRNA or 100% of ACT DMSO for Fabp4 mRNA levels. *p<05, **p<.01, ****p<.0001

### 5-aza-dC partially restores Thy1 cell surface expression in 3T3-L1 cells when exposed to adipogenic cocktail

We next examined Thy1 cell surface expression on 3T3-L1 cells, since Thy1 is a known cell marker on pre-adipocytes and is readily detected via flow cytometry and immunofluorescence. Cells were treated for 7 days daily with either DMSO or 5-aza-dC, while ACT samples were given the adipogenic cocktail starting at day 3, as previously described. The representative histogram in Figure 3A shows cells treated with ACT shift out of the Thy1+ gate into Thy1-during differentiation, while cells treated with ACT and 5-aza-dC mostly remain in the Thy1+ gate. As expected, pre-adipocytes treated with ACT expressed significantly less surface Thy1 than cells with media alone, as evidenced by a lower mean fluorescence intensity (MFI) (Fig 3B). However, pre-adipocytes cultured with ACT and 5-aza-dC showed a significant increase in Thy1 MFI compared to ACT alone, which occurred in tandem with an increase in the percentage of Thy1-positive cells (Fig 3B-C). Using immunofluorescence, we confirmed that Thy1 expression was sustained in ACT-treated pre-adipocytes also treated with 5-aza-dC, while Fabp4 expression decreased compared to cells treated with ACT alone (Fig. 3D).

**Figure 3:**
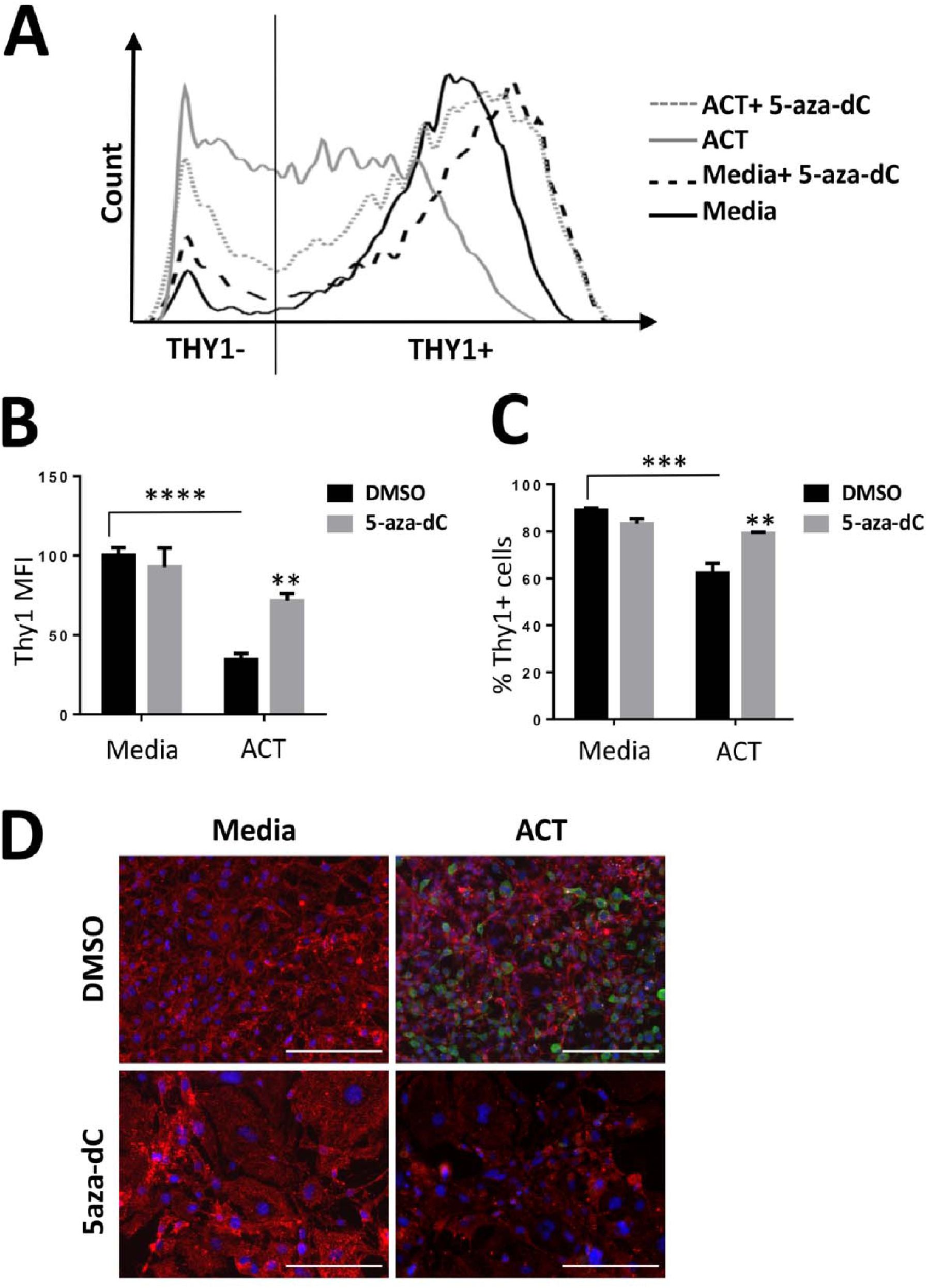
Thy1 surface protein levels attenuated by 5-aza-dC. 3T3-L1 cells were treated daily with either DMSO or 5-aza-dC for 7 days with the introduction of the adipogenic cocktail (ACT) starting at day 3. A) Representative histogram of Thy1 surface levels measured by flow cytometry using a CD90.2-PE conjugated antibody. B) ACT samples had a decrease in Thy1 mean fluorescence intensity (MFI) that was partially attenuated by 5-aza-dC treatment. C) ACT samples had a decrease in Thy1+ cells compared to Media alone, while ACT cells treated with 5-aza-dC had an increase in the Thy1+ population. D) Immunofluorescent images show decreased staining of Thy1 (Red) with ACT compared to pre-adipocytes (media alone) and increased Fabp4 (Green). Thy1 staining was sustained when ACT samples were treated with 5-aza-dC, along with a decreased staining of Fabp4. Cell nuclei are stained with DAPI and depicted in blue. Scale bars in white represent 200μm. **p<.01, ***p<.001, ****p<.0001

### Thy1(CD90) gene expression is regulated by DNA methylation during adipogenesis

We went on to examine *Thy1* DNA methylation, which typically occurs in cytosine (CG) rich regions and is commonly associated with gene silencing^34,35^. As the transcriptional activation of the *Thy1* gene involves both the promoter and intron 1 in some cell types^21,23^, and previous publications have referred to intron 1 as part of the promoter^17,19-23^, we focused on a CpG rich region within intron 1 that we termed Thy1-CGI1 (Fig 4A). Using a pyrosequencer, we analyzed methylation levels of 5 consecutive CpG sites within Thy1-CGI1. 3T3-L1 pre-adipocytes were treated as described previously. We found that Thy1-CGI1 is hypermethylated during differentiation comparing the average DNA methylation of ACT-treated samples to those treated with media alone. Treatment with 5-aza-dC resulted in reduced methylation of these CpG sites in the *Thy1* gene in both media alone and ACT-treated samples (Fig. 4B). Furthermore, individual CpG positions within Thy1-CGI1 showed an increase in DNA methylation when treated with the adipogenic cocktail versus media alone, with a significant increase at CpG position 2 (Fig 4C). Methylation decreased across all 5 CpG sites when treated with 5-aza-dC, whether the cells were treated with media alone or with ACT (Fig 4D-E). Four of the five CpG sites examined had significantly reduced methylation levels when cells were treated with ACT and 5-aza-dC. Our data indicates that this region is methylation sensitive and can influence *Thy1* expression during adipogenesis.

**Figure 4:**
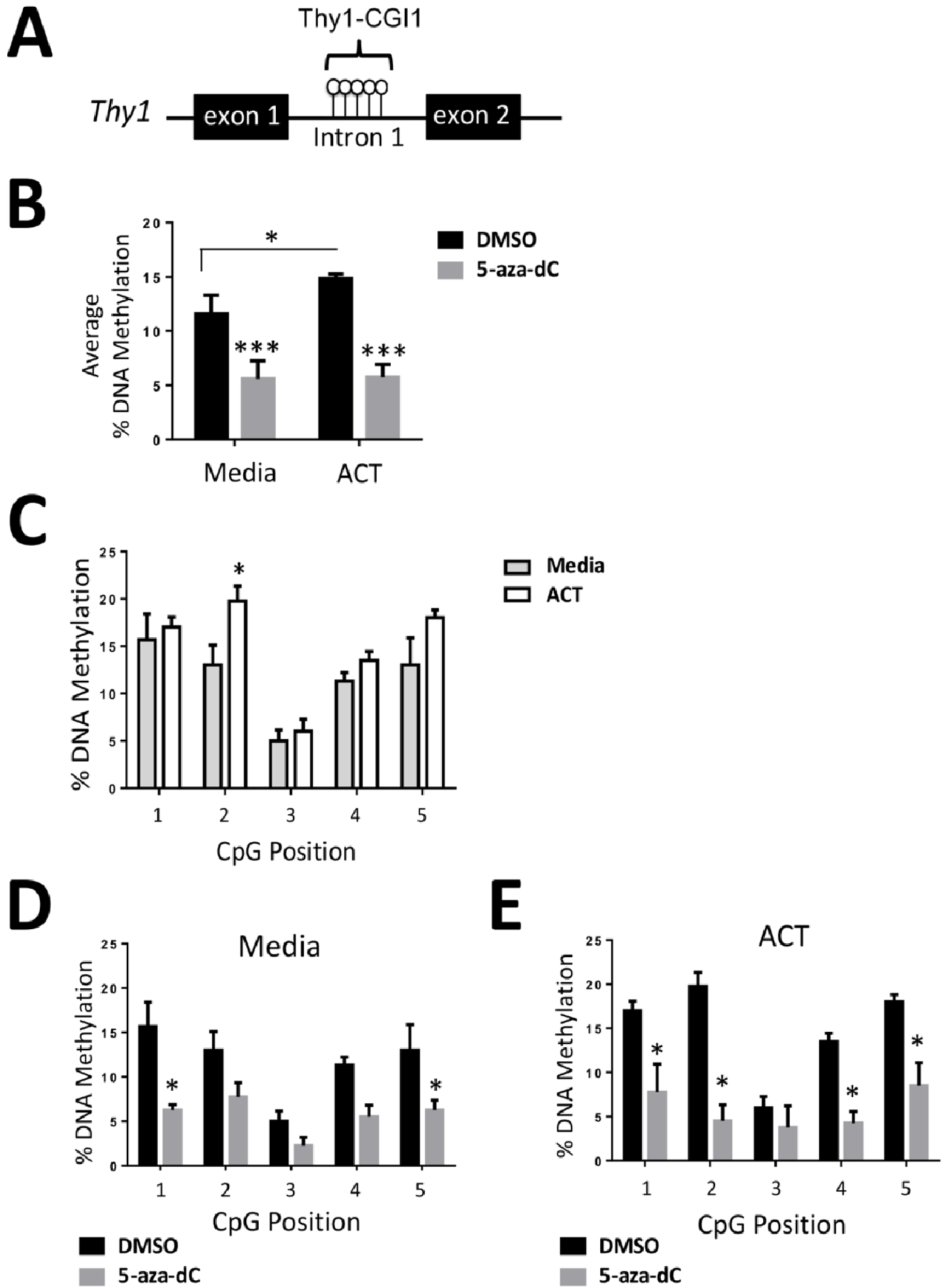
*Thy1* is hypermethylated during adipogenesis and demethylated by 5-aza-dC. 3T3-L1 cells were treated daily with either DMSO or 5-aza-dC for 7 days with the introduction of the adipogenic cocktail (ACT) starting at day 3. A) Schematic of *Thy1* gene showing Thy1-CGI1 and CpG sites between exon 1 and 2. B) Average percent DNA methylation of Thy1-CGI1 showed an increase in methylation in ACT cells compared to media alone, while treatment with the methylation inhibitor, 5-aza-dC resulted in a decrease of overall methylation in both groups. C) Individual CpG position sites were measured across Thy1-CGI1 and showed increases in methylation with ACT compared to media alone. D-E) Both media alone and ACT groups had decreases in methylation in all individual CpG sites when treated 5-aza-dC. * p<.05, ***p<.001

To test the methylation status of *Thy1* over time during adipocyte differentiation, we examined methylation levels at these five sites at days 0 (prior to ACT), 2, 4, and 6. Cells were pretreated with DMSO or given 5-aza-dC for 24 h, then either harvested at day 0 or given ACT medium and with continued treatment with DMSO (vehicle) or 5-aza-dC daily. Overall, average DNA methylation percentages increased in a time-dependent manner when exposed to the cocktail, with the highest degree of methylation occurring on day 6 (Fig 5A). Day 0 samples showed lower methylation levels compared to other time points in which cells had been exposed to ACT. Furthermore, there were no significant changes at any of the 5 CpG sites when given a single dose of 5-aza-dC (Fig. 5B). However, day 6 samples showed a significant two-fold increase in methylation compared to day 0, along with a significant decrease in methylation when cells were exposed daily to 5-aza-dC at all five CpG sites (Fig. 5C). This implies that during the normal adipogenic process, CpG sites at the *Thy1* locus become hypermethylated, which may blunt *Thy1* expression and allow for differentiation to occur.

**Figure 5:**
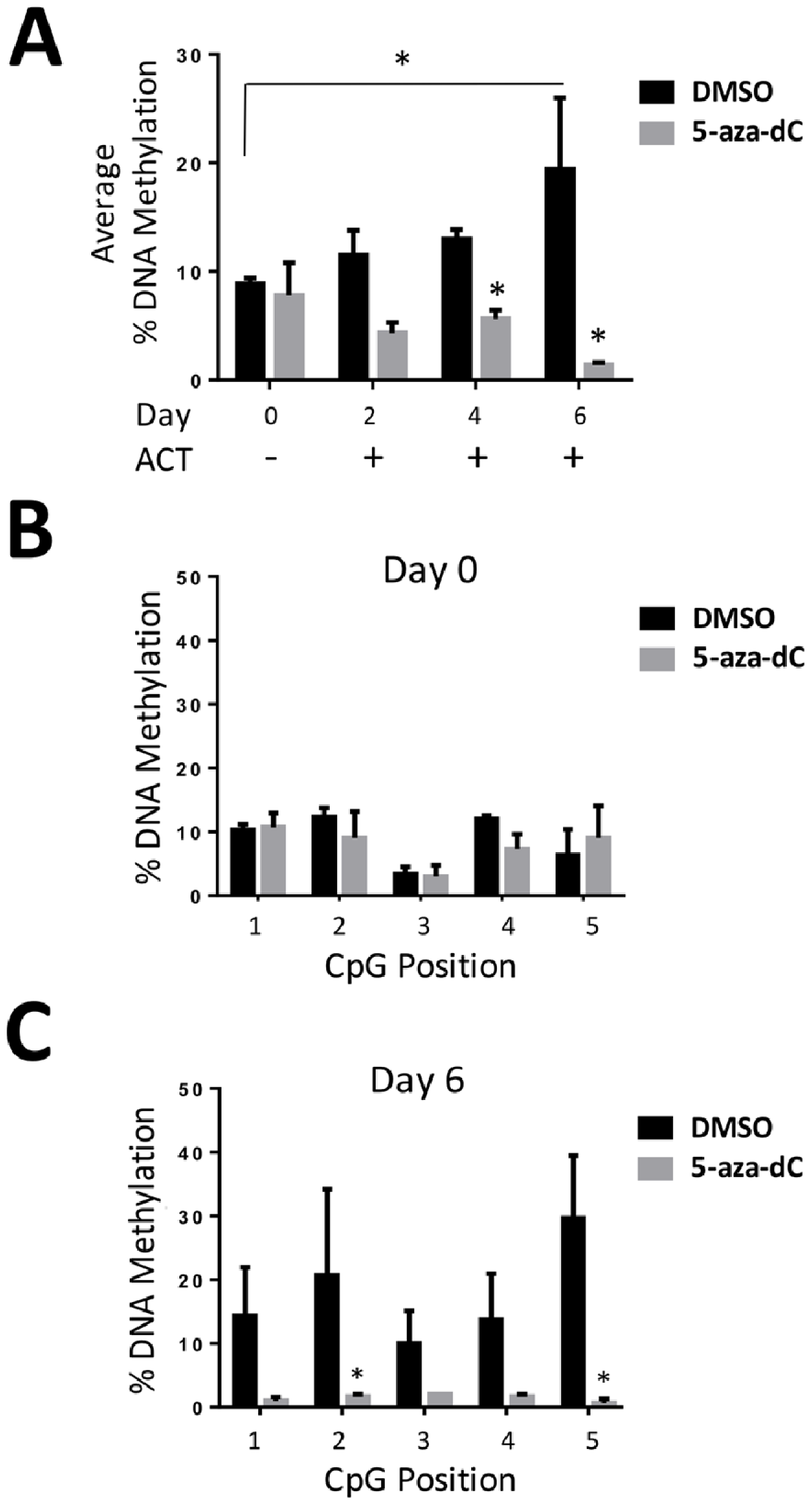
*Thy1* is hypermethylated in a time dependent manner during adipocyte differentiation and attenuated by 5-aza-dC. 3T3-L1 cells were pretreated for 24 h with either DMSO or 5-aza-dC, along with continuous daily treatment and then harvested corresponding to days with adipogenic cocktail (ACT) (Day 0= no ACT). A) Average percent DNA methylation of Thy1-CGI1 showed an increase in methylation in normal adipogenesis differentiation process, while treatment with the methylation inhibitor, 5-aza-dC resulted in a decrease in overall methylation. B-C) Individual CpG position sites were measured across Thy1-CGI1 and day 0 showed no relevant changes, while day 6 had a notable decrease in methylation when treated with 5-aza-dC. *= p<.05

## Discussion

Excessive adipogenesis can lead to weight gain and obesity, which affects over 700 million people worldwide. The consequences of obesity can be dire, including the development of cardiovascular or liver disease, diabetes, and other comorbidities, which result in significant morbidity and mortality. Therefore, understanding the adipogenic pathway(s) and molecular changes that foster adipocyte differentiation will elucidate the mechanisms contributing to the pathogenesis of obesity. New understanding should lead to better solutions for this growing problem, as lifestyle changes (e.g. improved diet and exercise) are often insufficient. Although genome-wide studies have shown that changes in histone methylation/acetylation occur during adipogenesis^25,36,37^, few specific adipogenesis-relevant genes have been identified as influenced by epigenetic changes (e.g. DNA methylation). In this study, we identify *Thy1* as a methylation sensitive gene and demonstrate that DNA methylation plays an active role in adipogenesis. Ultimately, we found that inhibiting DNA methylation blunts adipogenesis and sustains Thy1 at levels that may retard or suppress adipogenesis.

Thy1 is a cell surface protein that is expressed on mouse thymocytes and on both mouse and human pre-adipocytes^38,39^. We have previously shown in mouse 3T3-L1 cells that Thy1 is down-regulated in a time dependent manner during adipocyte differentiation, while cells overexpressing Thy1 cells no longer differentiate, even when given an adipogenic cocktail, which typically causes 3T3-L1 cells to differentiate into adipocytes after 6-8 days of exposure. However, here we show that 5-aza-dC blunts adipogenesis, even in the presence of ACT. These findings correlate with reduced levels of Fabp4 expression at both the protein and mRNA levels, along with partially attenuated levels of Thy1 when exposed to a global methylation inhibitor. Fabp4 levels inversely correlate with Thy1 levels; Fabp4 expression increases while Thy1 is suppressed during adipogenesis^14^. Therefore, blunted levels of Fabp4 are likely another contributing factor that impedes adipogenesis. We also demonstrated that during adipogenesis, Thy1-CGI1 methylation increases, which likely contributes to reduced *Thy1* expression. While we saw significant changes at CpG position 2 in ACT-treated samples relative to untreated (media alone) cells (Fig. 4C), we observed similar trends for the other CpG positions. These results are consistent with previous studies showing alterations in methylation status at CpG sites can cause significant changes in gene expression^40,41^. Treatment with the methylation inhibitor, 5-aza-dC, resulted in hypomethylation of Thy1-CGI1, which in part may contribute to the attenuation of *Thy1* expression. Since 5-aza-dC is a global DNA methylation inhibitor, it can affect other genes involved in adipogenesis in addition to *Thy1*. Our data suggest that DNA methylation is a necessary regulatory mechanism for pre-adipocytes to differentiate into fat cells, consistent with recent reports implying DNA methylation is involved in lineage-specific adipocyte development^24,42^.

Adipogenesis is a complex process that is controlled by many factors, which include epigenetic and post-transcriptional modifications. Previous studies have established that the expression/activity of microRNAs, small RNAs ~20-22 bp in length that bind to and block the transcription of specific targeted genes, play a role adipogenesis^43^. Furthermore, in obesity, it has been shown that there are significant changes in the expression of microRNAs (miR), such as an increase in miR-103 levels^44,45^. We have recently shown that miR-103 levels increase during adipogenesis and can bind the 3’ UTR of Thy1 to blunt its expression^15^. Here, we confirmed that the levels of miR-103 increase during adipogenesis, and that these levels were unchanged by treatment with 5-aza-dC, suggesting that the regulatory effects of *Thy1* methylation are separate from changes in microRNA expression (Figure S3). This represents another potential facet of *Thy1* regulation. Therefore, several, possibly overlapping mechanisms are involved in the adipogenic pathway. This has important implications for developing therapeutic interventions to combat adipogenesis and obesity; more than one aspect of this regulatory mechanism may need to be targeted to cause a significant effect.

Stem cells, such as, mesenchymal stem cells (MSCs) are distinguished and defined by various cell surface markers, including Thy1^46^. Mesenchymal stem cells can also differentiate into osteocytes and chondrocytes^39^, where Thy1 is expressed heterogeneously in each subset^47^. Recent studies have shown 5-aza-dC prevents adipogenesis and promotes osteoblastogenesis through the activation of Wnt10a^30^. Wnt10a is known to be upregulated and essential for bone formation^48-50^. However, Thy1 also plays a critical role^51,52^. It was recently shown that Thy1 is upregulated during osteoblastogenesis and that Thy1^-^ cells (knockdown and knockout) cannot differentiate into osteoblasts^53^. However, changes in *Thy1* DNA methylation during osteoblast formation have not been analyzed. In our present study, we show that Thy1 levels are sustained, while FABP4 levels are lowered in mMSCs treated with 5-aza-dC in the presence of ACT. While many epigenetic factors aid in stem cell maintenance,^54^ changes in the methylation status of genes, such as *Thy1*, may alter stem cell state. Since *Thy1* is highly expressed in MSCs and pre-adipocytes, maintaining *Thy1* expression may be the key to remaining a precursor cell. However, further investigation is needed. While we saw an increase in *Thy1* methylation in mouse pre-adipocyte cells, a crucial next step would be to determine basal *Thy1* expression levels and the methylation status of MSCs derived from adipose tissues of obese and non-obese individuals. Testing the ability of these cells to differentiate into fat cells may correlate with their *Thy1* expression profiles. Such investigations are likely to provide additional evidence of Thy1’s critical role in adipogenesis and would further underline the importance of our findings.

While we show that 5-aza-dC treatment decreased methylation in the Thy1-CGI1 region and blunted adipogenesis, it is possible that hypomethylating the *Thy1* gene at the same time could affect other cell types, such as, fibroblasts (involved in fibrosis); it has been established that *Thy1* expression is up-regulated during and involved in myofibroblast differentiation^55^. Future studies could examine other CpG islands in the traditional promoter region and downstream regions to investigate whether methylation of these sites is also essential for *Thy1’s* involvement in adipogenesis, fibrosis, and other functions. While the *Thy1* gene may be a key target for methylation during adipogenesis, there are undoubtedly other genes regulated by methylation during differentiation. Further investigation is necessary to identify other essential genes, which is fundamental to understanding the adipogenic pathway.

In summary, our work shows for the first time that *Thy1* has increased DNA methylation during adipogenesis. We demonstrate herein that inhibiting DNA methylation attenuates the loss of *Thy1* when cells are stimulated to differentiate into adipocytes. Blocking methylation leads to sustained *Thy1* expression and prevents adipogenesis. These studies further highlight the role of genomic methylation and *Thy1’s* involvement in adipogenesis, which suggests these pathways may be dysregulated in metabolic diseases in which adipogenesis is elevated, such as obesity.

## Materials and Methods

### Chemicals

5-aza-2′-deoxycytidine (5-aza-dC), 3-isobutyl-1-methylxanthine (IBMX), dexamethasone, and human recombinant insulin were all purchased from Sigma-Aldrich (St. Louis, MO).

### Cell Culture

All cells were incubated at 37°C with 7% humidified CO2. 3T3-L1 cells were maintained in 10% calf serum supplemented with DMEM media. C57BL/6 Mouse Bone Marrow Mesenchymal Stem Cells were purchased from Cell Biologics (Chicago, IL) and maintained in 10% mesenchymal stem cell-qualified fetal bovine serum in supplemented MEM media from Thermo Fisher. Cells were plated at 60% confluence and treated at 80% confluency. To induce adipogenesis, media containing an adipogenic cocktail (ACT) was added to confluent cells, which consists of 0.5 mM IBMX, 0.5 μM dexamethasone, and 2 μg/ml insulin. Fresh ACT was added every 2 days. To inhibit methylation, cells were treated daily with 0.5uM 5-aza-dC or DMSO as a control. Cells were then harvested on days indicated per experiment.

### Quantitative real-time PCR (qPCR) detection of mRNA

RNA was extracted with a Qiagen miRNeasy Kit and quantified using a NanoDrop 1000 spectrophotometer (Thermo Scientific, Wilmington, DE). A BioRad iScript reverse transcription kit was used to make cDNA from 150 ng RNA. RT-qPCR assays were then performed using SsoAdvanced Universal SYBR Green Supermix (Bio-Rad), according to the manufacturer’s instructions. All genes of interest were normalized to 18S rRNA, and the relative percentages were normalized to 100% to media + DMSO for Thy1 mRNA or 100% to ACT + DMSO for Fabp4 mRNA levels. Primer sequences were as follows Thy1: Fwd 5’-CCTTACCCTAGCCAACTTCAC and Rv 5’-AGGATGTGTTCTGAACCAGC; Fabp4: Fwd 5’-ATGTGTGATGCCTTTGTGGGAAC and Rv5’-TCATGTTGGGCTTGGCCATG; 18s rRNA: Fwd 5’-GTAACCCGTTGAACCCCATT and Rv 5’-CCATCCAATCGGTAGTAGCG.

### Western blot analysis

Cells were lysed with 60 mM Tris, 2% SDS, and protease inhibitor cocktail (Sigma-Aldrich). Ten μg of protein was loaded per lane and run on SDS-PAGE gels. Protein gels were transferred to 0.45 um Immobilon-PVDF membranes (Millipore, Temecula, CA) and blocked with 5% BSA in 0.1% Tween 20 in PBS. Primary antibodies, sheep anti-mouse Thy1 (R&D), rabbit anti-mouse Fabp4 (Cell Signaling), and rabbit anti-mouse β-tubulin (Cell Signaling) were diluted 1:5000, 1:500, and 1:5000, respectively, and incubated for 1 h. Membranes were washed in 0.1% Tween 20 in PBS then incubated in anti-sheep or anti-rabbit HRP-conjugated secondary antibodies at 1:5000 or 1:20,000 dilution, respectively. Protein was visualized using Immobilon Western chemiluminescent horseradish peroxidase substrate (Millipore). MagicMark XP protein standard protocol used for ladder (Novex). Blots were developed by X-ray film. All blots are provided as uncropped images in the supplementary data.

### Flow cytometry

Cells were trypsinized and washed in PBS, then fixed with 2% PFA and blocked with 1:50 human Fc receptor blocker (Miltenyi Biotech Inc., San Diego, CA) in PBS. The cells were then incubated with anti-mouse Thy1.2-PE conjugated antibody, 1:500, (BD Biosciences, San Jose, CA) for 1 h on ice. Cells were washed and resuspended in PBS. Cells were analyzed on a LSR II flow cytometer running FACSDIVA software (BD Biosciences). Analysis of fluorescence data was performed using FlowJo software v10.1 (FlowJo, LLC, Ashland, Oregon).

### Immunofluorescent staining

Cells treated in 12-well plates were washed with 1X PBS and fixed with 2% PFA for 10 min and washed three times with PBS. Cells were blocked in 1% BSA and 0.1% Triton X-100 in PBS with normal donkey serum (Jackson Immunoresearch) and Fc-blocker 1:50 (BD Biosciences). The primary antibodies used were Thy1.2-PE conjugated antibody, (BD Biosciences, San Jose, CA) and Fabp4 (Cell Signaling), which were diluted 1:500 in 1% BSA and incubated for 2 h at room temperature in the dark. After removal of primary antibody and three washes, secondary antibody (donkey anti-rabbit AF647) was applied at a 1:2000 dilution for an hour. Cells were then washed and visualized on an EVOS-FL Cell Imaging System (Thermo Fisher).

### DNA extraction and Bisulfite conversion

Genomic DNA was isolated from cells using a DNeasy DNA extraction kit (Qiagen, Valencia, CA) and quantified using the NanoDrop 1000 spectrophotometer (Thermo Scientific, Wilmington, DE). 1000 ng of genomic DNA was then bisulfite converted using an Epitect Plus Bisulfite Conversion Kit (Qiagen) to be analyzed by a pyrosequencer.

### Pyrosequencing assays

Bisulfite-treated DNA was amplified using the PyroMark PCR kit (Qiagen, Valencia, CA) with the conditions of, 95°C for 5 min, 45 cycles of (95°C for 30 s, annealing temperature of 58°C for 30 s, 72°C for 30 s), 72°C for 10 min. The PCR product sizes were then verified with electrophoresis on a 2% agarose gel. Ten µl of the biotinylated PCR products were mixed with 1 μl streptavidin-coated Sepharose beads, 40 μl PyroMark binding buffer (Qiagen), and 29 μl RNAase-free water for a total volume of 80 μl. This mixture was then run on a PyroMark Vacuum Workstation (Qiagen). The purified PCR products were then added to the annealing buffer, which contained the corresponding sequencing primer. After annealing, the plate was loaded into the PyroMark Q96 MD instrument (Qiagen). PyroMark-CpG software automatically generates a dispensation order of dNTPs and control dispensations, based on the sequence to analyze. Controls are included in the dispensation order to check the performance of the reactions. All runs also included a no template control. We analyzed the data with the PyroMark software for quantification of % DNA CpG methylation.

### Pyrosequencing Primers

Methylation levels were measured in the first CpG island of intron 1, which is part of the promoter^17,20-23^ of mouse *Thy1* (chr9:44,043,384-44,048,579; GRCm38/mm10) (94bp-349bp) using the pyrosequencing assay. Gene-specific primers for *Thy1* were designed using the Pyro-Mark assay design software, version 2.0 (Qiagen, Valencia, CA). The program automatically generated primer sets that included both PCR and sequencing primers, based on selected target sequences. One of the primers was biotinylated to enable immobilization to streptavidin-coated beads. The sequences were as follows: Forward primer: 5’-TTTAGTTATAGTTTTGGGAAAGGATAT Reverse Biotinylated primer: 5’-CCACCTCCTCCCTCTATT Sequencing primer: 5’-ATAGGGAFTTTTTATAT

### Statistical analysis

All values are presented as mean ± SEM. Experiments were conducted in triplicate at separate times. Two-way analysis of variance (ANOVA) were used for statistical analysis using GraphPad Prism6. P-values < 0.05 were considered significant.

## Acknowledgements

This project is funded by NIH grants F31ES027767, TL1-TR000096, ES001247, and ES0023032.

## Authors Contributions

E.F. performed all experiments. E.F. C.W. M.S. R.P. aided in experimental design and data analysis. E.F. prepared figures and wrote main manuscript. All authors aided in editing the manuscript. All authors have reviewed the manuscript.

## Additional Information

Competing Interests: The authors declare no competing interests.

## References

1. Afshin A, Forouzanfar MH, Reitsma MB, et al. Health Effects of Overweight and Obesity in 195 Countries over 25 Years. The New England journal of medicine. 2017;377(1):13–27.

2. Flegal KM, Kruszon-Moran D, Carroll MD, Fryar CD, Ogden CL. Trends in Obesity Among Adults in the United States, 2005 to 2014. Jama. 2016;315(21):2284–2291.

3. Ahima RS. Connecting obesity, aging and diabetes. Nature medicine. 2009;15(9):996–997.

4. Smyth S, Heron A. Diabetes and obesity: the twin epidemics. Nature medicine. 2006;12(1):75–80.

5. Heymsfield SB, Wadden TA. Mechanisms, Pathophysiology, and Management of Obesity. The New England journal of medicine. 2017;376(3):254–266.

6. Cawley J, Meyerhoefer C. The medical care costs of obesity: an instrumental variables approach. Journal of health economics. 2012;31(1):219–230.

7. Jo J, Gavrilova O, Pack S, et al. Hypertrophy and/or Hyperplasia: Dynamics of Adipose Tissue Growth. PLoS computational biology. 2009;5(3):e1000324.

8. Joe AW, Yi L, Even Y, Vogl AW, Rossi FM. Depot-specific differences in adipogenic progenitor abundance and proliferative response to high-fat diet. Stem cells (Dayton, Ohio). 2009;27(10):2563–2570.

9. Jing K, Heo JY, Song KS, et al. Expression regulation and function of Pref-1 during adipogenesis of human mesenchymal stem cells (MSCs). Biochimica et biophysica acta. 2009;1791(8):816–826.

10. Hudak CS, Sul HS. Pref-1, a gatekeeper of adipogenesis. Frontiers in endocrinology. 2013;4:79.

11. Kim CY, Kim GN, Wiacek JL, Chen CY, Kim KH. Selenate inhibits adipogenesis through induction of transforming growth factor-beta1 (TGF-beta1) signaling. Biochemical and biophysical research communications. 2012;426(4):551–557.

12. Kumar A, Ruan M, Clifton K, Syed F, Khosla S, Oursler MJ. TGF-beta mediates suppression of adipogenesis by estradiol through connective tissue growth factor induction. Endocrinology. 2012;153(1):254–263.

13. Cawthorn WP, Bree AJ, Yao Y, et al. Wnt6, Wnt10a and Wnt10b inhibit adipogenesis and stimulate osteoblastogenesis through a beta-catenin-dependent mechanism. Bone. 2012;50(2):477–489.

14. Woeller CF, O'Loughlin CW, Pollock SJ, Thatcher TH, Feldon SE, Phipps RP. Thy1 (CD90) controls adipogenesis by regulating activity of the Src family kinase, Fyn. FASEB journal : official publication of the Federation of American Societies for Experimental Biology. 2015;29(3):920–931.

15. Woeller CF, Flores E, Pollock SJ, Phipps RP. Editor’s Highlight: Thy1 (CD90) Expression is Reduced by the Environmental Chemical Tetrabromobisphenol-A to Promote Adipogenesis Through Induction of microRNA-103. Toxicological sciences : an official journal of the Society of Toxicology. 2017;157(2):305–319.

16. Robinson CM, Neary R, Levendale A, Watson CJ, Baugh JA. Hypoxia-induced DNA hypermethylation in human pulmonary fibroblasts is associated with Thy-1 promoter methylation and the development of a pro-fibrotic phenotype. Respiratory research. 2012;13:74.

17. Kolsto AB, Kollias G, Giguere V, Isobe KI, Prydz H, Grosveld F. The maintenance of methylation423 free islands in transgenic mice. Nucleic acids research. 1986;14(24):9667–9678.

18. Mann V, Szyf M, Razin A, Chriqui-Zeira E, Kedar E. Characterization of a tumorigenic murine T425 lymphoid-cell line spontaneously derived from an IL-2-dependent T-cell line. International journal of cancer. 1986;37(5):781–786.

19. Neveu WA, Mills ST, Staitieh BS, Sueblinvong V. TGF-beta1 epigenetically modifies Thy-1 expression in primary lung fibroblasts. American journal of physiology Cell physiology. 2015;309(9):C616–626.

20. Sanders YY, Pardo A, Selman M, et al. Thy-1 promoter hypermethylation: a novel epigenetic pathogenic mechanism in pulmonary fibrosis. American journal of respiratory cell and molecular biology. 2008;39(5):610–618.

21. Giguere V, Isobe K, Grosveld F. Structure of the murine Thy-1 gene. The EMBO journal. 1985;4(8):2017–2024.

22. Gundersen G, Kolsto AB, Larsen F, Prydz H. Tissue-specific methylation of a CpG island in transgenic mice. Gene. 1992;113(2):207–214.

23. Spanopoulou E, Giguere V, Grosveld F. The functional domains of the murine Thy-1 gene promoter. Molecular and cellular biology. 1991;11(4):2216–2228.

24. Lim YC, Chia SY, Jin S, Han W, Ding C, Sun L. Dynamic DNA methylation landscape defines brown and white cell specificity during adipogenesis. Molecular metabolism. 2016;5(10):1033–1041.

25. Broholm C, Olsson AH, Perfilyev A, et al. Human adipogenesis is associated with genome-wide DNA methylation and gene-expression changes. Epigenomics. 2016;8(12):1601–1617.

26. Guo W, Chen J, Yang Y, Zhu J, Wu J. Epigenetic programming of Dnmt3a mediated by AP2alpha is required for granting preadipocyte the ability to differentiate. Cell death & disease. 2016;7(12):e2496.

27. Sakamoto H, Kogo Y, Ohgane J, et al. Sequential changes in genome-wide DNA methylation status during adipocyte differentiation. Biochemical and biophysical research communications. 2008;366(2):360–366.

28. Yang X, Wu R, Shan W, Yu L, Xue B, Shi H. DNA Methylation Biphasically Regulates 3T3-L1 Preadipocyte Differentiation. Molecular endocrinology (Baltimore, Md). 2016;30(6):677–687.

29. Zhao QH, Wang SG, Liu SX, et al. PPARgamma forms a bridge between DNA methylation and histone acetylation at the C/EBPalpha gene promoter to regulate the balance between osteogenesis and adipogenesis of bone marrow stromal cells. The FEBS journal. 2013;280(22):5801–5814.

30. Chen YS, Wu R, Yang X, et al. Inhibiting DNA methylation switches adipogenesis to osteoblastogenesis by activating Wnt10a. Scientific reports. 2016;6:25283.

31. Xu S, De Becker A, Van Camp B, Vanderkerken K, Van Riet I. An improved harvest and in vitro expansion protocol for murine bone marrow-derived mesenchymal stem cells. Journal of biomedicine & biotechnology. 2010;2010:105940.

32. Qian SW, Li X, Zhang YY, et al. Characterization of adipocyte differentiation from human mesenchymal stem cells in bone marrow. BMC developmental biology. 2010;10:47.

33. Lehmann GM, Woeller CF, Pollock SJ, et al. Novel anti-adipogenic activity produced by human fibroblasts. American journal of physiology Cell physiology. 2010;299(3):C672–681.

34. Suzuki MM, Bird A. DNA methylation landscapes: provocative insights from epigenomics. Nature reviews Genetics. 2008;9(6):465–476.

35. Hsieh CL. Dependence of transcriptional repression on CpG methylation density. Molecular and cellular biology. 1994;14(8):5487–5494.

36. Keller M, Hopp L, Liu X, et al. Genome-wide DNA promoter methylation and transcriptome analysis in human adipose tissue unravels novel candidate genes for obesity. Molecular metabolism. 2017;6(1):86–100.

37. Mikkelsen TS, Xu Z, Zhang X, et al. Comparative epigenomic analysis of murine and human adipogenesis. Cell. 2010;143(1):156–169.

38. Haeryfar SM, Hoskin DW. Thy-1: more than a mouse pan-T cell marker. Journal of immunology (Baltimore, Md : 1950). 2004;173(6):3581–3588.

39. Orbay H, Tobita M, Mizuno H. Mesenchymal stem cells isolated from adipose and other tissues: basic biological properties and clinical applications. Stem cells international. 2012;2012:461718.

40. Attwood JT, Yung RL, Richardson BC. DNA methylation and the regulation of gene transcription. Cellular and molecular life sciences : CMLS. 2002;59(2):241–257.

41. Baylin SB, Herman JG, Graff JR, Vertino PM, Issa JP. Alterations in DNA methylation: a fundamental aspect of neoplasia. Advances in cancer research. 1998;72:141–196.

42. Bowers RR, Kim JW, Otto TC, Lane MD. Stable stem cell commitment to the adipocyte lineage by inhibition of DNA methylation: role of the BMP-4 gene. Proceedings of the National Academy of Sciences of the United States of America. 2006;103(35):13022–13027.

43. Du J, Cheng X, Shen L, et al. Methylation of miR-145a-5p promoter mediates adipocytes differentiation. Biochemical and biophysical research communications. 2016;475(1):140–148.

44. Li M, Liu Z, Zhang Z, Liu G, Sun S, Sun C. miR-103 promotes 3T3-L1 cell adipogenesis through AKT/mTOR signal pathway with its target being MEF2D. Biological chemistry. 2015;396(3):235–244.

45. Perri R, Nares S, Zhang S, Barros SP, Offenbacher S. MicroRNA modulation in obesity and periodontitis. Journal of dental research. 2012;91(1):33–38.

46. Alipour R, Sadeghi F, Hashemi-Beni B, et al. Phenotypic characterizations and comparison of adult dental stem cells with adipose-derived stem cells. International journal of preventive medicine. 2010;1(3):164–171.

47. Yoshimura H, Muneta T, Nimura A, Yokoyama A, Koga H, Sekiya I. Comparison of rat mesenchymal stem cells derived from bone marrow, synovium, periosteum, adipose tissue, and muscle. Cell and tissue research. 2007;327(3):449–462.

48. Kasaai B, Moffatt P, Al-Salmi L, Lauzier D, Lessard L, Hamdy RC. Spatial and temporal localization of WNT signaling proteins in a mouse model of distraction osteogenesis. The journal of histochemistry and cytochemistry : official journal of the Histochemistry Society. 2012;60(3):219–228.

49. Rudnicki MA, Williams BO. Wnt signaling in bone and muscle. Bone. 2015;80:60–66.

50. Yin X, Wang X, Hu X, Chen Y, Zeng K, Zhang H. ERbeta induces the differentiation of cultured osteoblasts by both Wnt/beta-catenin signaling pathway and estrogen signaling pathways. Experimental cell research. 2015;335(1):107–114.

51. Chung MT, Liu C, Hyun JS, et al. CD90 (Thy-1)-positive selection enhances osteogenic capacity of human adipose-derived stromal cells. Tissue engineering Part A. 2013;19(7–8):989–997.

52. Bearden RN, Huggins SS, Cummings KJ, Smith R, Gregory CA, Saunders WB. In-vitro characterization of canine multipotent stromal cells isolated from synovium, bone marrow, and adipose tissue: a donor-matched comparative study. Stem cell research & therapy. 2017;8(1):218.

53. Paine A, Woeller CF, Zhang H, et al. Thy1 is a positive regulator of osteoblast differentiation and modulates bone homeostasis in obese mice. FASEB journal : official publication of the Federation of American Societies for Experimental Biology. 2018:fj201701379R.

54. Fernandez-Arroyo S, Cuyas E, Bosch-Barrera J, Alarcon T, Joven J, Menendez JA. Activation of the methylation cycle in cells reprogrammed into a stem cell-like state. Oncoscience. 2015;2(12):958–967.

55. Hansen TC, Woeller CF, Lacy SH, Koltz PF, Langstein HN, Phipps RP. Thy1 (CD90) Expression Is Elevated in Radiation-Induced Periprosthetic Capsular Contracture: Implication for Novel Therapeutics. Plastic and reconstructive surgery. 2017;140(2):316–326.

